# A method for predicting linear and conformational B-cell epitopes in an antigen from its primary sequence

**DOI:** 10.1101/2023.01.18.524531

**Authors:** Nishant Kumar, Sadhana Tripathi, Neelam Sharma, Sumeet Patiyal, Naorem Leimarembi Devi, Gajendra P. S. Raghava

## Abstract

B-cell is an essential component of the immune system that plays a vital role in providing the immune response against any pathogenic infection by producing antibodies. Existing methods either predict linear or conformational B-cell epitopes in an antigen. In this study, a single method was developed for predicting both types (linear/conformational) of B-cell epitopes. The dataset used in this study contains 3875 B-cell epitopes and 3996 non-B-cell epitopes, where B-cell epitopes consist of both linear and conformational B-cell epitopes. Our primary analysis indicates that certain residues (like Asp, Glu, Lys, Asn) are more prominent in B-cell epitopes. We developed machine-learning based methods using different types of sequence composition and achieved the highest AUC of 0.80 using dipeptide composition. In addition, models were developed on selected features, but no further improvement was observed. Our similarity-based method implemented using BLAST shows a high probability of correct prediction with poor sensitivity. Finally, we came up with a hybrid model that combine alignment free (dipeptide based random forest model) and alignment-based (BLAST based similarity) model. Our hybrid model attained maximum AUC 0.83 with MCC 0.49 on the independent dataset. Our hybrid model performs better than existing methods on an independent dataset used in this study. All models trained and tested on 80% data using cross-validation technique and final model was evaluated on 20% data called independent or validation dataset. A webserver and standalone package named “CLBTope” has been developed for predicting, designing, and scanning B-cell epitopes in an antigen sequence (https://webs.iiitd.edu.in/raghava/clbtope/).

## Introduction

The immune system is an interactive and interconnected network of cells, tissues, and organs that work together to defend the host against foreign invaders [1]. It recognizes and responds to a wide range of disease-causing pathogens such as parasites, fungus, bacteria and viruses. The immune system can be subdivided into innate and adaptive immunity [2]. Innate immune response provides first line of protection against invasive pathogens and activates adaptive immune system. It is a non-specific, immediate reaction which does not produce any immunological memory [3]. On the other hand, adaptive immunity provides antigen-specific defense and generates lifelong immunological memory [4]. It is further composed of B and T-lymphocytes which identify antigens with distinct specificity. B cells stimulate humoral immunity, whereas T lymphocytes stimulate cell-mediated immunity [5].

Antibodies secreted by B-cells play an important role in the immune response. It specifically binds to antigens at the antigenic determinant site, also known as epitopes. An epitope is made up of proteins that are present on the surface of the antigen. It interacts with the antibody through the B-cell receptor and evokes either cellular or humoral immunological response [6]. Based on the structural characteristics of epitopes and their interaction with antibodies, they can be classified into linear (or continuous) or conformational (discontinuous) epitopes [7]. Linear epitopes are a stretch of continuous residues of amino acids required for binding, whereas conformational epitopes contain amino acids that are far apart from each other but come in close proximity due to polypeptide folding [8]. B-cell epitopes provide a great deal of potential for immunology-related applications, such as the development of epitope-based vaccines, therapeutic antibodies, and disease diagnosis [9, 10]. Although experimental techniques for identification B-cell epitopes are more accurate but very expensive, time-consuming and labour-intensive when compared to computational methods [11].

To predict B-cell epitopes, several computational methods have been developed in the past [12]. These method can be mainly divided into three categories; conformational, linear and linearized conformational B-cell epitopes. A region/segment on surface of a protein structure recognize by B-cell receptors is called conformational or discontinuous B-cell epitope. Numerous methods have been developed to predict conformational B-cell epitopes from protein structures like CEP, DiscoTope, SEPPA, Epitope3D and SEMA [13–17]. These conformational methods need tertiary structure of a protein to identify conformational B-cell epitopes in a protein. Thus, prediction of tertiary structure of a protein is one of the major bottleneck in implementation of conformational methods. A single continuous stretch of amino acids within a protein sequence that can reacted with anti-protein antibodies is predicted by the majority of modern methods, which are known as linear/continuous B-cell epitope prediction methods. Initially, linear methods were developed using physicochemical properties [18, 19], most of them have very poor performance [20]. This leads to development machine learning based methods that include ABCpred, BepiPred and BCPREDS [21–23]. These methods were developed using small datasets mainly extracted from BCIPEP, AntiJen 2.0 and HIVDB. Next generation of methods extract dataset from IEDB and used machine learning and deep learning for methods for predicting continuous B-cell epitopes [24–26]. In addition, methodologies have been designed to predict conformational epitopes mainly antibody-interacting residues in an antigen from it is primary sequence [27, 28].

Recently, linearized conformational epitopes were identified from antibody-antigen structures for developing epitope prediction [29]. A strategy to predict both linear and conformational B-cell epitopes has been developed in this study. Dataset used in this study contain linear and linearized conformational B-cell epitopes. Firstly, we developed machine learning based models using composition of epitope/non-epitope sequences. Secondly, we develop an alignment-based approach using BLAST, a peptide is assigned epitope if it has high similarity with known B-cell epitopes. It was observed that both techniques have their own limitation. Finally, using both alignment-based and alignment-free methods, we created a hybrid model. For the benefit of the scientific community, we developed a standalone software and web server that allow user to predict B-cell epitopes.

## Material and Methods

### Data Acquisition

We derived most of the datasets for this study from following datasets BCETD_555_, ILED_2195_, and IDED_1246_ obtained from Ras-Carmona *et. al*. [30]. BCETD_555_ is a non-redundant dataset containing linearized conformational 555 B-cell epitopes and 555 non B-cell epitopes retrieved from antibody-antigen structure complexes. ILED_2195_ contains 2195 linear B-cell epitopes and 2195 non-epitopes extracted from Immune Epitope Database (IEDB). Another conformational dataset IDED_1246_ obtain from IEDB, contains 1246 discontinuous B-cell epitopes and 1246 non-epitopes. We merged all three datasets and created a new dataset that contains 7992 epitopes (3996 B-cell epitopes and 3996 non-B-cell epitopes). We have also taken into consideration the length of peptide sequences while distributing the dataset. To enhance the quality of the dataset, we have removed the peptides comprising unnatural amino acids ‘BJOUXZ’ and duplicate sequences. Hence, the final dataset consists of 7871 epitopes (3875 B-cell and 3996 non-B-cell epitopes). Our B-cell epitopes have both types of B-cell epitopes, conformational and linear/continuous.

### Composition Analysis

Amino acid composition (AAC) represents the percentage frequency of 20 amino acids within peptide/protein sequences. It is a feature vector of 20 elements in which each component of the sequence represents a fraction of a specific type of amino acid residue in the sequence. It can be calculated by the following equation

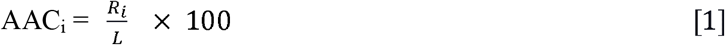

Where AAC_i_ is the percent composition of amino acid i; R_i_ is the number of residues of type i, and L is the total number of residues in the peptide [31–34].

### Sequence Logo

Sequence logos are generated to display sequence conservation patterns that provide a more detailed and accurate explanation of sequence similarity than consensus sequences. We used a web-based application, “weblogo” to generate sequence logos. It generates a graphical representation of a stack of amino acids measured in bits. Each stack’s overall height reveals sequence conversation at that place. Furthermore, the relative frequency of the relevant amino or nucleic acid at that place in the sequences is indicated by the height of the symbols within the stack [35].

### Feature Generation

To develop a powerful predictor, it is required to convert input peptide sequences into a set of numerical vectors/features representing the properties of the peptide sequence [36]. The extracted features must preserve the peptide’s sequence information to the greatest extent possible and reflect intrinsic correlation with the peptide classification [33]. For extracting features such as AAC, DPC, ATC, PCP, RRI (See Table 1) from peptide sequence, we have utilized the ‘Pfeature’ standalone package [37]. With this tool thousands of features/descriptors of protein and peptide sequences were calculated [38, 39]. In this study, we have used a composition-based feature module and generated a vector of 780 features, as shown in Table 1.

**Table 1.**
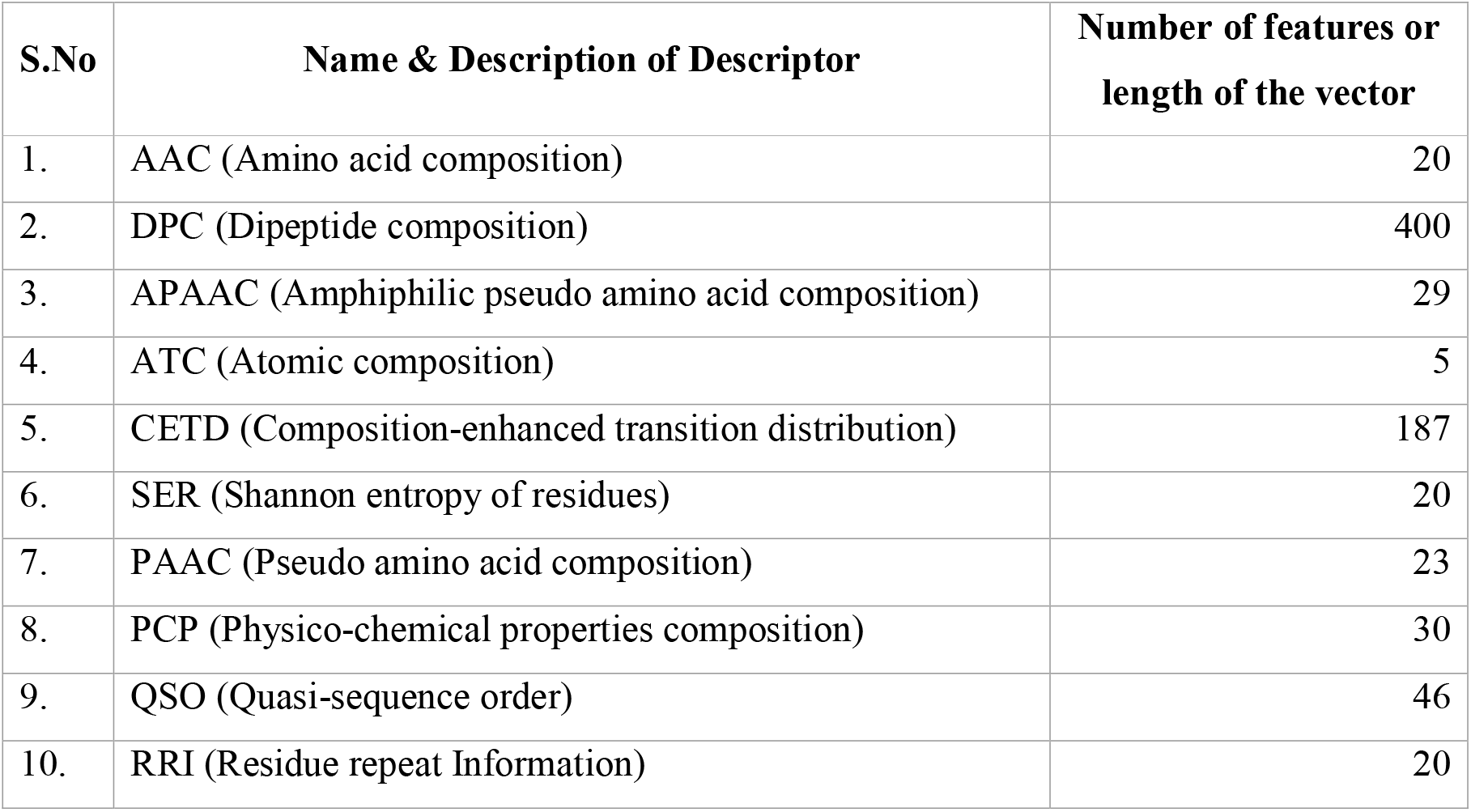
List of descriptors with brief description and number of features; all features computed using Pfeature.

| S.No | Name & Description of Descriptor | Number of features or length of the vector |
| --- | --- | --- |
| 1. | AAC (Amino acid composition) | 20 |
| 2. | DPC (Dipeptide composition) | 400 |
| 3. | APAAC (Amphiphilic pseudo amino acid composition) | 29 |
| 4. | ATC (Atomic composition) | 5 |
| 5. | CETD (Composition-enhanced transition distribution) | 187 |
| 6. | SER (Shannon entropy of residues) | 20 |
| 7. | PAAC (Pseudo amino acid composition) | 23 |
| 8. | PCP (Physico-chemical properties composition) | 30 |
| 9. | QSO (Quasi-sequence order) | 46 |
| 10. | RRI (Residue repeat Information) | 20 |

### Motif based features

A motif is a repeated pattern of sequences in a set of peptides or proteins. Identifying high-class motifs for positive dataset characterization involves comparing positive dataset motifs with those of negative datasets [32].

### Feature Selection

Feature selection is a crucial stage in classification. In most datasets, not all the features contribute to determining the endpoint; some of the features are redundant or noisy. “Pfeature” generates a large vector of features. It is a major challenge to identify the relationship between the features of the data. The ultimate objective of feature selection is to choose a pertinent subset of features from the initial feature set that can reduce the likelihood of over-fitting and boost effectiveness by simplifying the model [34, 36, 39–41]. A primary concern of this study is the ability to generate better models as fast as possible.

### Machine learning

Machine learning (ML) algorithms are powerful and potent techniques that are capable of transforming data into reliable decisions. We implement several machine learning algorithms [41] to develop a powerful predictor that can efficiently and accurately predict the B-cell epitopes. Our study adopt supervised classification algorithms, where each ML algorithm has a unique set of hyperparameters that should be tuned and modified to identify the optimal combination and, as a result, the best prediction and solution to the problem [34]. One major task is choosing an appropriate classification algorithm [36]. We have used the following ML algorithms: Decision Tree (DT), Random Forest (RF), Gaussian Naive Bayes (GNB), Logistic Regression (LR), XGBoost (XGB), k-nearest neighbors (KNNs), and Extra-Trees (ET) Classifier [38]. We must carefully train the ML algorithm to classify accurately on unknown datasets [34].

### Model’s Evaluation

For the evaluation of our model, we have implemented a five-fold cross-validation algorithm. As per the standard protocols, this technique splits data randomly into k-folds. We implemented cross-validation technique with k=5, then we trained our model on 4 folds, and rest one fold was used to test the model and the same cycle was repeated five times. In the results, we have shown the average scores of all repetitions [38, 41].

### Similarity Search

For protein /peptide annotation a well-known similarity search based technique used called “BLAST”. In this technique, all peptides whose functions are known are aligned with the query peptide. The query peptide is annotated based on its alignment score with known peptides. In our study, we implemented the BLAST-based technique blastp (BLAST+2.7.1), a peptide-peptide BLAST for the prediction of B-cell epitopes and non B-cell epitopes [42–45]. BLAST formatted database were constructed using the training dataset against which the query sequences (sequences in the test set) were hit at various e-values that ranges from 1e-6 to 1e+3. On the basis of the top hits, the query sequence was classified as positive if the top hit was positive and vice versa if the top hit was negative.

### Evaluation parameters

Several parameters are taken into account to measure the quality or predictive performance of ML algorithm [32]. The parameters considered to evaluate model performance are threshold dependent and threshold-independent. Threshold dependent parameters include: Sensitivity, Specificity, Accuracy, and Matthew’s correlation coefficient (MCC), whereas Area Under the Receiver Operating Characteristic (AUROC) is considered a threshold-independent parameter These performance evaluation criteria have been widely applied to evaluate the model’s effectiveness and they are well-defined in the literature [31, 38, 41, 46]. The measurements are defined as follows:

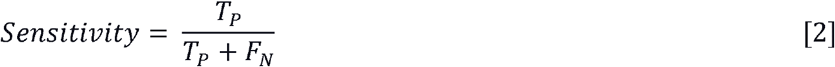

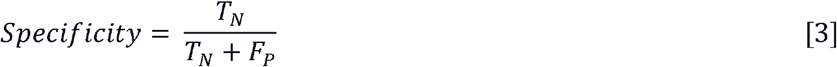

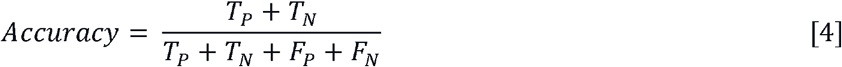

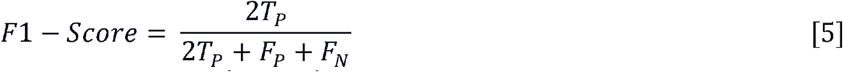

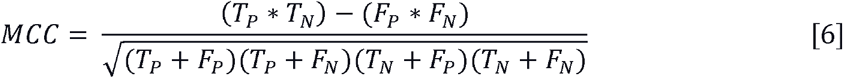

Where, T_N_, T_P_, F_N_, and F_P_ stands for true negative, true positive, false negative, and false positive respectively.

### Web server implementation

We have built a web server called “CLBTope” to predict B-cell epitopes (https://webs.iiitd.edu.in/raghava/clbtope). The user-friendly and device-compatible front end of CLBTope was created using HTML5, CSS, and PHP scripts. Users can submit a sequence in FASTA format through the online server’s Predict, Design, Protein Scan, Motif Scan, and Blast Scan modules.

## Results

### Compositional analysis

AAC provides the occurrence frequency of a given peptide, which allows to easily distinguish between the B-cell epitopes from non B-cell epitopes. Here, we compute the average composition of B-cell epitopes, non B-cell epitopes, and general proteome, as shown in Figure 1. The amino-acid residues lysine (K), aspartic acid (D), glutamic acid (E), asparagine (N), arginine (R), proline (P), and glutamine (Q) are most abundant in the positive dataset, whereas residues alanine (A), cysteine (C), phenylalanine (F), isoleucine (I), valine (V), and leucine (L) are highly conserved in the negative dataset.

**Figure 1:**
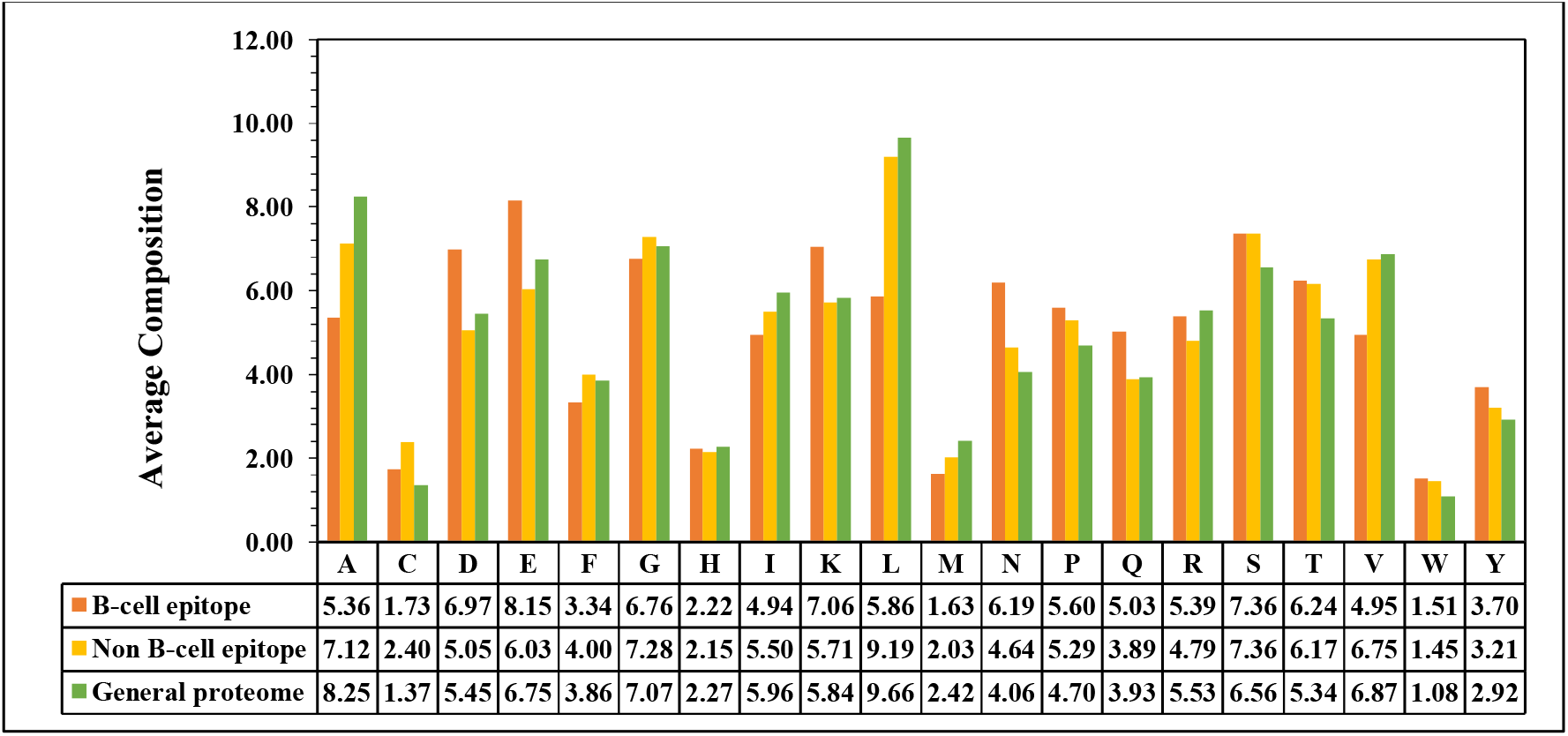
Percent average amino acid composition of B-cell epitopes, non B-cell epitopes, and general proteome.

### Positional analysis

In this study, we have built/constructed a sequence logo to understand and inspect the inclination of a particular residue at a specific position in B-cell epitopes. As depicted in Figure 2, the hydrophilic residue–glutamic acid (E) is a highly abundant and conserved residue at almost all the positions, whereas, serine (S) is more preserved at 3^rd^, 6^th^, 7^th^, and 21^st^ position; however, lysine (K) dominates the 18^th^ position.

**Figure 2:**
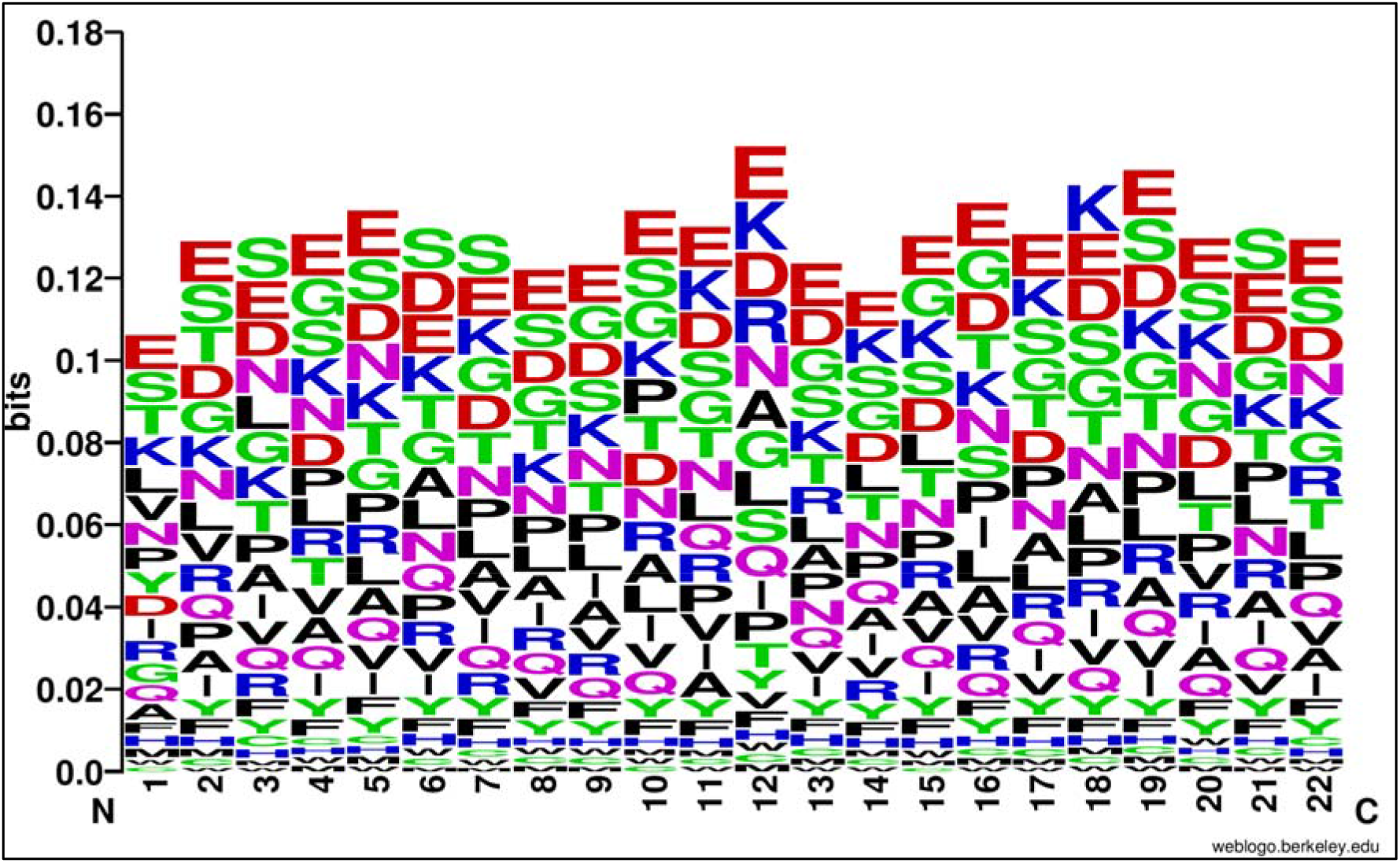
Sequence logo of B-cell epitopes, glutamic acid (E) residue is dominant at most of the positions.

### B-cell Epitope Prediction Models

For the purpose of developing antibodies with desired specificity and designing vaccines, B-cell epitope identification is of practical interest [29]. We have developed a B-cell epitope prediction model as a classification problem for ML that can accurately distinguish B-cell epitopes from non B-cell epitopes. In this study, we construct a dataset from “BCEPS” that includes 3875 B-cell epitopes, which we call a positive dataset, and 3996 non B-cell epitopes, called a negative dataset. The system architecture of the method is provided in Figure 3.

**Figure 3:**
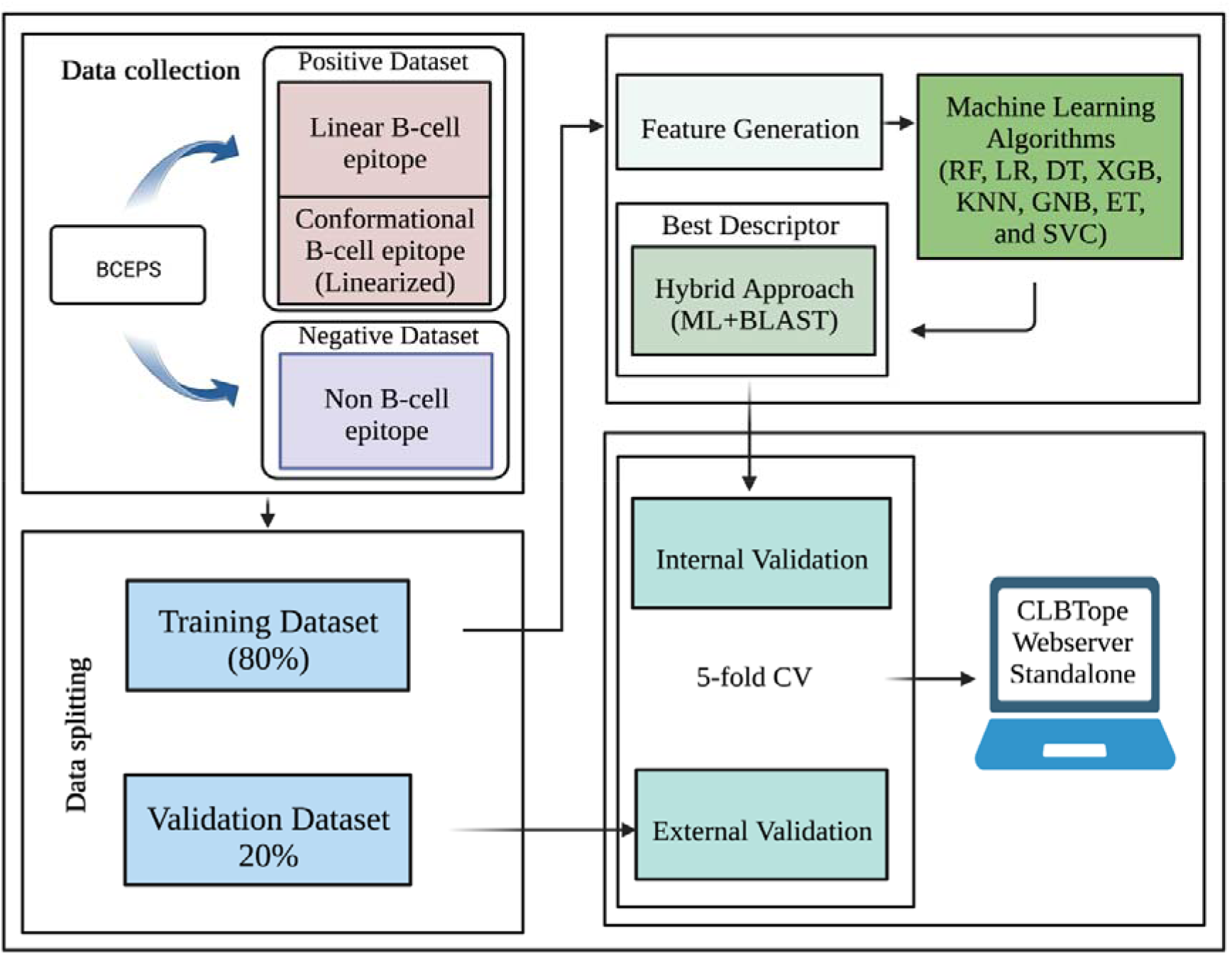
Complete workflow of the study.

We specially applied various ML algorithms such as DT, RF, LR, XGB, KNN, GNB, and ET to build a model. One of the crucial phase is the selection of the most appropriate ML algorithm for classification. Firstly, we develop prediction models on 10 different composition-based features extracted using the “Pfeature” tool. It was observed that RF (Random forest) performs best among other ML-algorithms and achieved almost similar performance on all the descriptors achieving AUROC above 0.72. In the 5-fold cross validation, our model achieved the highest AUROC of 0.802 on DPC-400 features on the validation dataset, which was significantly better than other descriptors shown in Table 2. In Supplementary Table S1, comprehensive results of several classifiers are provided.

**Table 2:** Performance of ML-based models developed on 10 different composition-based features using RF algorithm.

| Name & Description of descriptor | Training |  | Validation |  |
| --- | --- | --- | --- | --- |
|  | AUROC | MCC | AUROC | MCC |
| DPC (Dipeptide composition) | 0.790 | 0.437 | 0.802 | 0.442 |
| PAAC (Pseudo amino acid composition) | 0.787 | 0.410 | 0.796 | 0.449 |
| SER (Shannon entropy of residues) | 0.787 | 0.412 | 0.794 | 0.449 |
| APAAC (Amphiphilic pseudo amino acid composition) | 0.787 | 0.417 | 0.793 | 0.435 |
| AAC (Amino acid composition) | 0.788 | 0.419 | 0.793 | 0.450 |
| CETD (Composition-enhanced transition distribution) | 0.782 | 0.428 | 0.792 | 0.416 |
| SPC (Shannon entropy of physico-chemical properties) | 0.763 | 0.377 | 0.763 | 0.388 |
| PCP (Physico-chemical properties composition) | 0.760 | 0.383 | 0.762 | 0.406 |
| QSO (Quasi-sequence order) | 0.763 | 0.387 | 0.761 | 0.389 |
| RRI (Residue repeat Information) | 0.719 | 0.309 | 0.726 | 0.319 |

#AUROC, Area Under Receiver Operating Curve; MCC, Matthews correlation coefficient

In our analysis, most descriptors have comparatively similar performance to DPC. Therefore, we concatenate all the descriptors [34, 36] that achieve AUROC above 0.760, producing a hybrid feature vector of 760, including the proportion of DPC-400. The hybrid feature vector includes DPC, PAAC, SER, APAAC, AAC, SPC, PCP, QSO, and CETD. Here, we observed that RF performs best as it achieves maximum performance with an AUROC 0.800 and has a balanced sensitivity and specificity (Shown in Table 3).

**Table 3:** Performance of machine-learning-based models developed on Hybrid features combination.

| Name | Training |  |  |  |  | Validation |  |  |  |  |
| --- | --- | --- | --- | --- | --- | --- | --- | --- | --- | --- |
|  | Sens | Spec | Acc | AUROC | MCC | Sens | Spec | Acc | AUROC | MCC |
| **DT** | 66.516 | 65.561 | 66.031 | 0.705 | 0.321 | 68.903 | 64.581 | 66.709 | 0.702 | 0.335 |
| **RF** | 72.129 | 71.723 | 71.923 | 0.793 | 0.438 | 70.839 | 73.091 | 71.982 | 0.800 | 0.439 |
| **LR** | 68.645 | 69.346 | 69.001 | 0.762 | 0.380 | 68.387 | 69.962 | 69.187 | 0.761 | 0.384 |
| **XGB** | 71.387 | 71.067 | 71.224 | 0.787 | 0.424 | 72.129 | 70.713 | 71.410 | 0.786 | 0.428 |
| **KN** | 69.484 | 70.472 | 69.986 | 0.773 | 0.400 | 69.935 | 70.839 | 70.394 | 0.774 | 0.408 |
| **GNB** | 68.419 | 69.096 | 68.763 | 0.736 | 0.375 | 67.097 | 70.338 | 68.742 | 0.741 | 0.375 |
| **ET** | 72.097 | 71.786 | 71.939 | 0.791 | 0.439 | 72.387 | 72.966 | 72.681 | 0.801 | 0.454 |

#Sens, Sensitivity; Spec, Specificity; Acc, Accuracy; AUROC, Area Under Receiver Operating Curve; MCC, Matthews correlation coefficient; DT, Decision Tree; RF, Random Forest; LR, Logistic Regression; XGB, XGBoost; KNNs, k-nearest neighbors; GNB, Gaussian Naive Bayes; ET, Extra-Trees Classification

### Performance of selected features

A large number of the feature includes unrelated features that consume a lot of computing time to train the model. The model is simplified when developed on extracted truly relevant features [33]. In this study, we adopt the mRMR algorithm (minimum redundancy-maximum relevance) [40] to remove unrelated features or interdependence between features. We analyzed the performance on the top-400, 200, and 100 selected set of hybrid features and observed that the maximum AUROC achieved is 0.79 among the above selected sets. Figure 4 shows the 5-fold cross validation prediction results on hybrid features with feature selection (mRMR) and without feature selection. Supplementary Table S2 includes all of the results.

**Figure 4:**
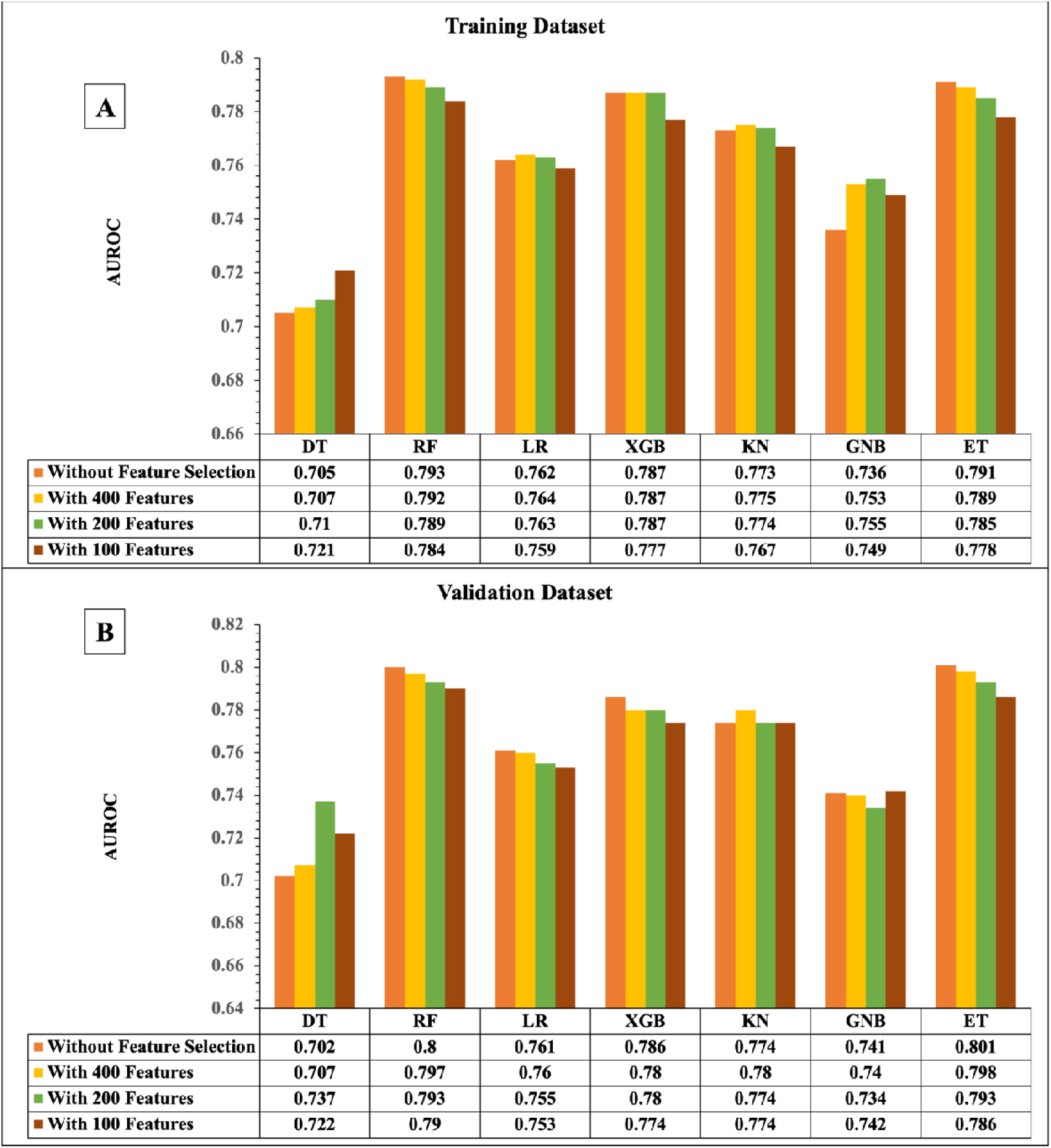
Performance of different classifiers with and without feature selection (A) Training dataset, (B) Validation dataset.

The performance on DPC composition (a vector of 400 features) is still superior to hybrid feature in terms of AUROC, shown in Table 4. Furthermore, we focus on DPC descriptor to improve the performance and selection of best features using the mRMR feature selection algorithm. We extract a unique set of features from the DPC descriptor and evaluate the performance on top-200, 150, and 100 selected features. It was shown that a DPC-400 features is superior to the performance of these selected sets of features. In Figure 5, different classifiers based on different feature sets for training and validation datasets are compared (in terms of AUROC). In Supplementary Table S3, we provide the complete results.

**Figure 5:**
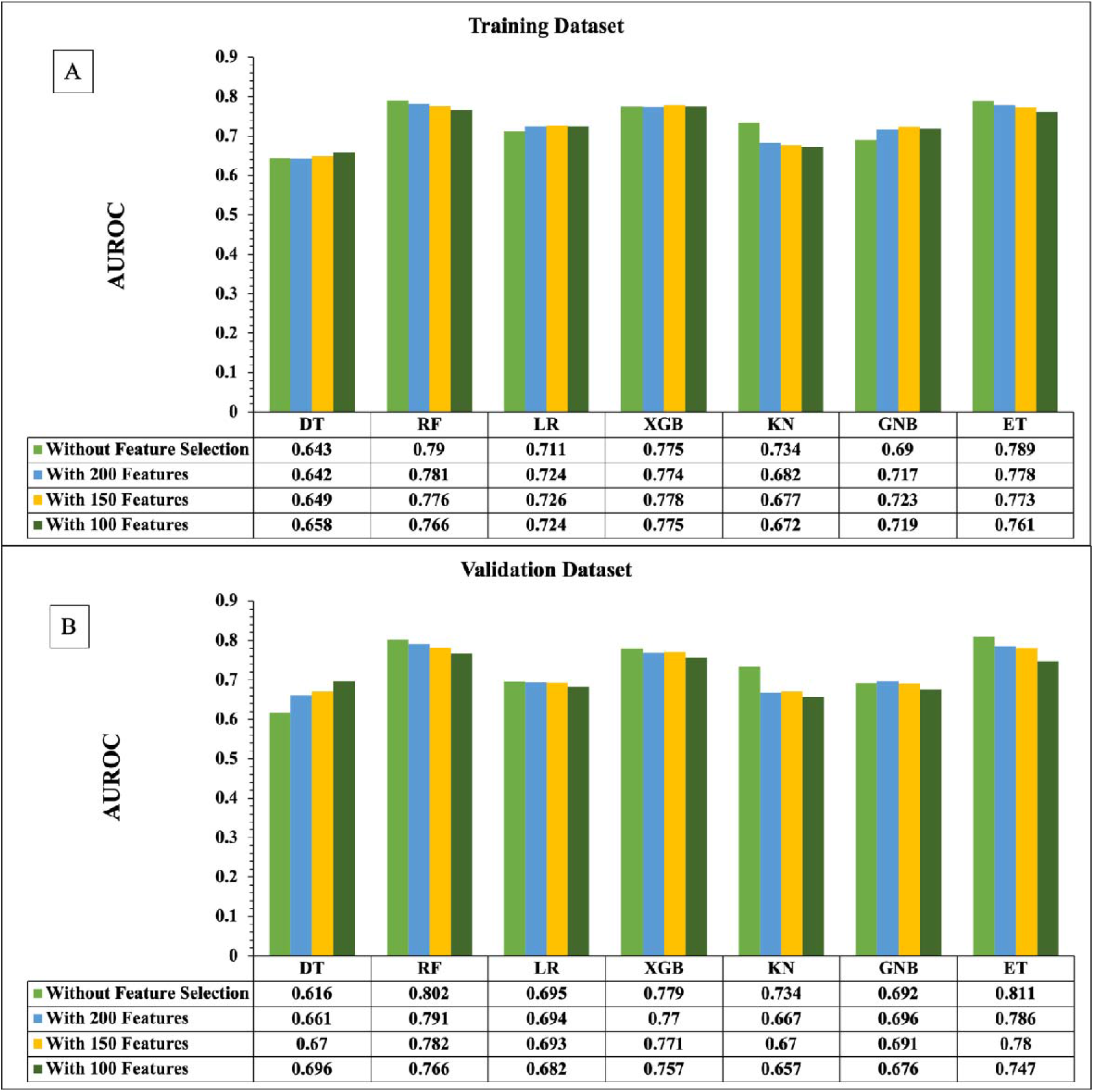
Performance of different classifiers on DPC-400, 200, 150, and 100 features (A) Training dataset, (B) Validation dataset.

**Table 4:** Performance of machine-learning-based models developed on DPC-400 features.

| Classifier | Training |  |  |  |  | Validation |  |  |  |  |
| --- | --- | --- | --- | --- | --- | --- | --- | --- | --- | --- |
|  | Sens | Spec | Acc | AUROC | MCC | Sens | Spec | Acc | AUROC | MCC |
| **DT** | 61.419 | 63.059 | 62.252 | 0.643 | 0.245 | 61.677 | 59.950 | 60.801 | 0.616 | 0.216 |
| **RF** | 71.613 | 72.537 | 72.082 | 0.790 | 0.442 | 72.000 | 72.340 | 72.173 | 0.802 | 0.443 |
| **LR** | 64.968 | 64.748 | 64.856 | 0.711 | 0.297 | 61.677 | 65.707 | 63.723 | 0.695 | 0.274 |
| **XGB** | 70.323 | 71.160 | 70.748 | 0.775 | 0.415 | 70.194 | 71.464 | 70.839 | 0.779 | 0.417 |
| **KN** | 67.484 | 66.250 | 66.857 | 0.734 | 0.337 | 68.000 | 63.579 | 65.756 | 0.734 | 0.316 |
| **GNB** | 63.935 | 63.935 | 63.935 | 0.690 | 0.279 | 65.161 | 64.581 | 64.867 | 0.692 | 0.297 |
| **ET** | 72.032 | 70.691 | 71.351 | 0.789 | 0.427 | 75.355 | 69.712 | 72.490 | 0.811 | 0.451 |

#Sens, Sensitivity; Spec, Specificity; Acc, Accuracy; AUROC, Area Under Receiver Operating Curve; MCC, Matthews correlation coefficient; DT, Decision Tree; RF, Random Forest; LR, Logistic Regression; XGB, XGBoost; KNNs, k-nearest neighbors; GNB, Gaussian Naive Bayes; ET, Extra-Trees Classification

### Similarity search approach

As BLAST is previously used for annotating and assigning functions to proteins based on similarity searches, we have developed a similarity search-based module to further improve the model performance [42, 47]. We implemented a similar concept (blastp) for annotating the given peptide as a B-cell epitope or non B-cell epitope. The query sequences (test set sequences) were searched against a local database that we created using the training dataset at e-values ranging from 1e^-6^ to 1e^+3^. Each query sequence has been categorized as B-cell epitope or non B-cell epitope based on top-hit. For instance, if the query sequence has the top-hit against a B-cell epitope, then the query peptide is assigned as a B-cell epitope, otherwise it is assigned as non B-cell epitope. As shown in Table 5, probability of correct prediction (chits) ranges from 1.94% - 70.32% on B-cell epitope dataset and 1.63% - 64.83% on non B-cell epitope dataset.

**Table 5:** The performance of BLAST-based search on the validation dataset.

| E-value | Positive Dataset |  |  | Negative Dataset |  |  |
| --- | --- | --- | --- | --- | --- | --- |
|  | Chits | Whits | No-hits | Chits | Whits | No-hits |
| **1.00E-06** | 15 (1.94%) | 1 (0.13%) | 759 (97.94%) | 13 (1.63%) | 1 (0.13%) | 785 (98.25%) |
| **1.00E-05** | 18 (2.32%) | 3 (0.39%) | 754 (97.29%) | 19 (2.38%) | 1 (0.13%) | 779 (97.5%) |
| **1.00E-04** | 34 (4.39%) | 9 (1.16%) | 732 (94.45%) | 27 (3.38%) | 9 (1.13%) | 763 (95.49%) |
| **1.00E-03** | 57 (7.35%) | 12 (1.55%) | 706 (91.10%) | 41 (5.13%) | 11 (1.38%) | 747 (93.49%) |
| **1.00E-02** | 85 (10.97%) | 17 (2.19%) | 673 (86.84%) | 59 (7.38%) | 19 (2.38%) | 721 (90.24%) |
| **1.00E-01** | 165 (21.29%) | 25 (3.23%) | 585 (75.48%) | 89 (11.14%) | 21 (2.63%) | 689 (86.23%) |
| **1.00E+00** | 282 (36.39%) | 30 (3.87%) | 463 (59.74%) | 118 (14.77%) | 25 (3.13%) | 656 (82.10%) |
| **1.00E+01** | 386 (49.81%) | 64 (8.26%) | 325 (41.94%) | 205 (25.66%) | 75 (9.39%) | 519 (64.96%) |
| **1.00E+02** | 518 (66.84%) | 205 (26.45%) | 52 (6.71%) | 467 (58.45%) | 265 (33.17%) | 67 (8.39%) |
| **2.00E+02** | 538 (69.42%) | 227 (29.29%) | 10 (1.29%) | 505 (63.20%) | 280 (35.04%) | 14 (1.75%) |
| **1.00E+03** | 545 (70.32%) | 229 (29.55%) | 1 (0.13%) | 518 (64.83%) | 281 (35.17%) | 0 (0.00%) |

\*Chits: correct hits; Whits: wrong hits

### Hybrid Methods

we combine two approaches to develop models called hybrid methods [44]. We integrate ML score with BLAST and MERCI score.

#### Hybrid1 approach

In this model, we incorporate BLAST similarity search score and machine learning prediction. The RF-based classifier performs best when utilising the DPC-400 feature vector, as shown in Tables 2 and 4. Therefore, our first step is to compute the ML prediction score using our best performing model. Secondly, we use BLAST at different evalues to classify a given peptide. Then, we assigned “+0.5” for a correct positive prediction (B-cell epitope),‘-0.5’ for a correct negative prediction (non B-cell epitope), and ‘0’ if hit is not found [42, 47, 48]. In order to predict B-cell epitope/non B-cell epitope, we integrate the BLAST score and ML-score for each query sequence to get the prediction performance. Finally, we developed a hybrid model and calculate the performance at different e-values (see Table 6). As shown in Table 6, at e-value ‘1’, we obtained a maximum AUROC of 0.829 with an accuracy of 74.460 on validation datasets.

**Table 6:** Model performance developed using the Hybrid method (BLAST and DPC-400) on training and validation dataset.

| E-Value | Training |  |  |  |  | Validation |  |  |  |  |
| --- | --- | --- | --- | --- | --- | --- | --- | --- | --- | --- |
|  | Sens | Spec | Acc | AUROC | MCC | Sens | Spec | Acc | AUROC | MCC |
| **1.00E-05** | 71.645 | 72.693 | 72.177 | 0.791 | 0.443 | 72.903 | 72.340 | 72.618 | 0.806 | 0.452 |
| **1.00E-04** | 71.677 | 72.599 | 72.145 | 0.790 | 0.443 | 72.903 | 71.965 | 72.427 | 0.801 | 0.449 |
| **1.00E-03** | 71.677 | 72.631 | 72.161 | 0.790 | 0.443 | 73.419 | 71.840 | 72.618 | 0.804 | 0.453 |
| **1.00E-02** | 71.839 | 72.693 | 72.273 | 0.791 | 0.445 | 72.387 | 73.467 | 72.935 | 0.803 | 0.459 |
| **1.00E-01** | 72.097 | 72.631 | 72.368 | 0.793 | 0.447 | 73.161 | 74.218 | 73.698 | 0.813 | 0.474 |
| **1.00E+00** | 72.484 | 72.693 | 72.590 | 0.796 | 0.452 | 74.323 | 74.593 | 74.460 | 0.829 | 0.489 |
| **1.00E+01** | 72.548 | 72.693 | 72.622 | 0.794 | 0.452 | 74.065 | 74.343 | 74.206 | 0.821 | 0.484 |

#Sens, Sensitivity; Spec, Specificity; Acc, Accuracy; AUROC, Area Under Receiver Operating Curve; MCC, Matthews correlation coefficient; RF, Random Forest

#### Hybrid2 approach

In this approach, we combine the DPC-400 ML-score and motif score. In order to find new sequences engaged in a biological process of interest, we used a technique called "MERCI" that locates motifs made up of particular amino acids and physico-chemical properties that can be used as discriminators. This tool includes an option to find gapped motifs and also introduces the two parameters FP and FN. The FP represents the minimal frequency threshold for the positive sequences, and FN represents the maximal frequency threshold for the negative sequences [49]. To identify the specific motifs in the positive dataset we have constructed a positive dataset known to be B-cell epitopes and a negative dataset of non B-cell epitopes. By using this dataset, this tool identifies top k motifs that occur most frequently in the positive dataset [49]. We started individually with simple 1-gap, 2-gap, and Koolman and Rohm classification without gaps. We also consider Koolman and Rohm classification with 1-gap and 2-gap. We integrate the DPC-400 ML score with Motif score to calculate the performance of the hybrid model. Finally, we achieved maximum AUROC with default parameters is 0.67 and AUROC with parameter FN is 0.71.

### Performance comparison with existing methods

We also performed the comparison with some existing B-cell prediction methods such as iBCEEL [50], ABCPred [21], LBTope [24], CBTope [27], BCEPred [20], and BepiPred3.0 [28], on validation dataset. Further, we obtain other performance measures including Sensitivity, Specificity, Accuracy, AUROC, and MCC, shown in Table 7.

**Table 7:** Comparative analysis of existing methods of B-cell prediction

| Name | Sens | Spec | Acc | AUROC | MCC |
| --- | --- | --- | --- | --- | --- |
| **iBCEEL** | 48.129 | 43.805 | 45.934 | 0.447 | 0.081 |
| **ABCPred** | 57.435 | 54.092 | 55.684 | 0.579 | 0.115 |
| **LBTope** | 55.742 | 55.945 | 55.845 | 0.583 | 0.117 |
| **CBTope** | 57.032 | 59.199 | 58.132 | 0.606 | 0.162 |
| **BCEPred** | 61.677 | 60.451 | 61.055 | 0.646 | 0.221 |
| **BepiPred3.0** | 63.484 | 62.703 | 63.088 | 0.696 | 0.262 |

#Sens, Sensitivity; Spec, Specificity; Acc, Accuracy; AUROC, Area Under Receiver Operating Curve; MCC, Matthews correlation coefficient

### Webserver Implementation

To provide better accessibility and navigation to scientific community, a user-friendly web server has been developed named “CLBTope” (https://webs.iiitd.edu.in/raghava/clbtope/) for the prediction of B-cell epitopes, either the epitope is linear or conformational (linearized). We implement our top-performing model in the webserver with the following modules: “Predict”, “Design”, “Scan”, “Motif Scan”, and “Blast Scan”. Users can classify submitted sequences as B-cell epitopes or non B-cell epitopes using the “Predict” module. By using the ‘Protein Scan’ module, users can scan or identify regions in an amino-acid sequence that correspond to B-cells or non-B-cells. Through the ‘Design’ module, users can generate all B-cell epitope analogs that can be created in the submitted sequence. The ‘Blast Scan’ module allows them to search the query sequence against the database of known B-cell epitopes. A query sequence is predicted as a B-cell epitope, or non B-cell epitope depending upon the match or hit in the database. If found matched or hit in the database, predicted as B-cell epitope; otherwise non B-cell epitope. A FASTA file format is also available for users to download the positive and negative datasets that we used in this study.

## Discussion and Conclusion

Immune responses protect us against foreign parasitic infection. Epitopes are antigenic determinant regions of antigen that are recognized by the host immune system, specifically by antibodies. Epitope based vaccine design is considered important to curb unforeseen pandemics and epidemics caused due to outbreak of infectious diseases. Epitope based peptide vaccine has an advantage over conventional vaccines as it facilitates the precise delivery of targeted vaccines. Prediction of B-cell epitope not only aids immunologists in designing epitope based peptide vaccines but also promotes the development of accurate diagnostic kits and treatment [51][52]. We have introduced a new method, “CLBTope” for the prediction of B-cell epitope and non B-cell epitope. Our model trained on the dataset with 3875 B-cell epitopes (positive dataset) and 3996 non B-cell epitopes (negative dataset) retrieved from BCEPS. We have used “Pfeature” for extracting the features from peptide sequences. Here, we compute the hybrid feature vector of 760 and apply following ML algorithms: DT, RF, LR, XGB, KNN, GNB, and ET. The hybrid vector was further reduced using the mRMR feature selection algorithm. Among all the models, the RF-based model outperforms and achieves the maximum AUROC of 0.802 on the DPC descriptor (a vector of 400 features). To further improve the model’s performance, we developed an RF-based hybrid model by combining the BLAST and DPC-400; the new method achieves the maximum AUROC 0.829 on the validation dataset. Building a reliable and accurate model for the prediction of B-cell epitope is important for vaccines that can be developed using targeted epitope regions and are capable enough to elicit host immune responses with minimal side effects. We anticipate that this method will significantly contribute to the scientific community working in this domain. We have provided a user-friendly webserver CLBTope (https://webs.iiitd.edu.in/raghava/clbtope/) for predicting, scanning, and designing B-cell epitopes.

## Funding Source

The current work has been supported by the Department of Biotechnology (DBT) funding agency.

## Conflict of interest

The authors declare no competing financial and non-financial interests.

## Authors’ contributions

GPSR collected the dataset. NK, SP, ST, and NS processed the dataset. SP, NK, ST, and GPSR implemented the algorithms and developed the prediction models. SP, NK, ST, NS, NLD, and GPSR analysed the results. NK and SP created the front-end and back-end of the webserver. NK, ST, SP, NLD, NS and GPSR penned the manuscript. GPSR conceived and coordinated the project. All authors have read and approved the final manuscript.

## Acknowledgements

Authors are thankful to the University Grants Commission (UGC), Department of BioTechnology (DBT), DBT-RA program in Biotechnology and Life Sciences, and Department of Science and Technology (DST-INSPIRE) for fellowships and financial support, and the Department of Computational Biology, IIITD New Delhi for infrastructure and facilities. We would like to acknowledge that Figures were created using BioRender.com.

## Data Availability Statement

All the datasets used in this study are available at the “CLBTope” web server, https://webs.iiitd.edu.in/raghava/clbtope/algo.php#dataset

## Notes

### Competing Interest Statement

The authors have declared no competing interest.

